# Model ensembles with different response variables for base and meta models: malaria disaggregation regression combining prevalence and incidence data

**DOI:** 10.1101/548719

**Authors:** Tim C. D. Lucas, Anita Nandi, Michele Nguyen, Susan Rumisha, Katherine E. Battle, Rosalind E. Howes, Chantal Hendriks, Andre Python, Penny Hancock, Ewan Cameron, Pete Gething, Daniel J. Weiss

## Abstract

Maps of infection risk are a vital tool for the elimination of malaria. Routine surveillance data of malaria case counts, often aggregated over administrative regions, is becoming more widely available and can better measure low malaria risk than prevalence surveys. However, aggregation of case counts over large, heterogeneous areas means that these data are often underpowered for learning relationships between the environment and malaria risk. A model that combines point surveys and aggregated surveillance data could have the benefits of both but must be able to account for the fact that these two data types are different malariometric units. Here, we train multiple machine learning models on point surveys and then combine the predictions from these with a geostatistical disaggregation model that uses routine surveillance data. We find that, in tests using data from Colombia and Madagascar, using a disaggregation regression model to combine predictions from machine learning models trained on point surveys improves model accuracy relative to using the environmental covariates directly.

## 1. Introduction

High-resolution maps of malaria risk are vital for elimination but mapping malaria in low burden countries presents new challenges as traditional mapping of prevalence from cluster-level surveys (Gething et al., 2011; Bhatt et al., 2017; Gething et al., 2012; Bhatt et al., 2015) is often not effective because, firstly, so few individuals are infected that most surveys will detect zero cases, and secondly, because of the lack of nationally representative prevalence surveys in low burden countries (Sturrock et al., 2016, 2014). Routine surveillance data of malaria case counts, often aggregated over administrative regions defined by geographic polygons, is becoming more reliable and more widely available (Sturrock et al., 2016) and recent work has focussed on methods for estimating high-resolution malaria risk from these data (Sturrock et al., 2014; Wilson and Wakefield, 2017; Law et al., 2018; Taylor et al., 2017; Li et al., 2012). However, the aggregation of cases over space means that the data may be relatively uninformative, especially if the case counts are aggregated over large or heterogeneous areas, because it is unclear where within the polygon, and in which environments, the cases occurred. This data is therefore often under-powered for fitting flexible, non-linear models as is required for accurate malaria maps (Bhatt et al., 2017, 2015). A model that combines point surveys and aggregated surveillance data, and therefore leverages the strength of both, has great potential.

One approach for combining these data is to use prevalence point-surveys to train a suite of machine learning models, and then use predictions from these models as covariates in a model trained on polygon-level incidence data. This process of stacking models has proven effective in many realms however typical stacking uses a single dataset on a consistent scale (Sill et al., 2009; Bhatt et al., 2017). Here we propose training the level zero machine learning models on point-level, binomial prevalence data and stacking these models with a polygon-level, Poisson incidence model.

## 2. Methodology

We used two data sources that reflect *Plasmodium falciparum* malaria transmission; point-prevalence surveys and polygon-level, aggregated incidence data. We selected Colombia and Madagascar as case examples as they both have fairly complete, publicly available, surveillance data at a finer geographical resolution than admin 1. The prevalence survey data were extracted from the Malaria Atlas Project prevalence survey database using only data from 1990 onwards (Bhatt et al., 2015; Guerra et al., 2007). For Colombia we used all points from South America (n = 522) while for Madagascar we used only Malagasy data (n = 1505). We chose these geographic regions based on a trade-off between wanting a large sample size but wanting data from geographically similar areas. The prevalence points were then standardised to an age range of 2–10 using the model from (Smith et al., 2007). The polygon incidence data were collected from government reports and standardised using methods defined in Cibulskis et al. (2011). This standardisation step accounts for missed cases due to lack of treatment seeking, missing case reports, and cases that sought medical attention outside the public health systems (Battle et al., 2016). For reports where cases were not reported at the species level, national estimates of the ratio between *P. falciparum* and *P. vivax* cases were used to calculate P. *falciparum* only cases. To minimise temporal effects we selected, for each country, one year of surveillance data. We used annual surveillance data from 2015 for Colombia (952 municipalities) and data from 2013 for Madagascar (110 districts) as these years had the most data in each case.

We considered an initial suite of environmental and anthropological covariates, at a resolution of approximately 5*×*5 kilometres that included the annual mean and log standard deviation of land surface temperature, enhanced vegetation index, malaria parasite temperature suitability index, elevation, tasseled cap brightness, tasseled cap wetness, log accessibility to cities, log night lights and proportion of urban land cover (Weiss et al., 2015). Tasseled cap brightness and urban land cover were subsequently removed as they were highly correlated with other variables. The covariates were standardised to have a mean of zero and a standard deviation of one. These covariates were used for both the machine learning models and the polygon-level models. Raster surfaces of population for the years 2005, 2010 and 2015, were created using data from WorldPop (Tatem, 2017) and from GPWv4 (NASA, 2018) where WorldPop did not have values. Population rasters for the remaining years were created by linear interpolation.

For each country we fitted five models via *caret* (Kuhn et al., 2017): elastic net (Zou and Hastie, 2012), Random Forest (Wright and Ziegler, 2015), projection pursuit regression (Friedman and Stuetzle, 1981), neural networks (Venables and Ripley, 2002) and boosted regression trees (Ridgeway et al., 2017). Our response variable was prevalence and we weighted the data by sample size (i.e. the number of people tested for malaria in each survey. For each model we ran five-fold cross-validation to select hyperparameters using random search for Random Forest and boosted regression trees and grid search for the other models. Predictions from these models were then made across Colombia and Madagascar respectively. These predictions were finally inverse logit transformed so that they are on the linear predictor scale of the top level model.

The top level model was a disaggregation regression model (Sturrock et al., 2014; Wilson and Wakefield, 2017; Law et al., 2018; Taylor et al., 2017; Li et al., 2012). This model is defined by a likelihood at the level of the polygon with covariates and a spatial random field at the pixel-level. Values at the polygon-level are given the subscript *a* while pixel level values are indexed with *b*.

The polygon case count data, *y*_*a*_ is given a Poisson likelihood

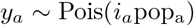

where *i*_*a*_ is the estimated polygon incidence rate and pop_a_ is the observed polygon population-at-risk. This polygon-level likelihood is linked to the pixel level prevalence

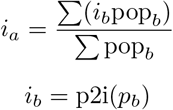

where p2i is from a model that was published previously (Cameron et al., 2015) which defines a function

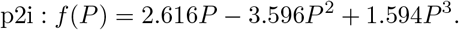

The fact that the model passes through prevalence space ensures that the predictions from the machine learning models can be appropriately scaled. The linear predictor of the model is related to prevalence by a typical logit link function and includes an intercept, *β*_0_, covariates, *X* with regression parameters *β*, a spatial, Gaussian, random field, *u*(*s, ρ, σ*_*u*_), and an *iid* random effect, *v*_*j*_ (*σ*_*v*_).

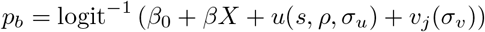

The Gaussian spatial effect has a Matérn covariance function and two hyperparameters: *ρ*, the nominal range (beyond which correlation is < 0.1) and *σ*_*u*_, the marginal standard deviation. The *iid* random effect models both missing covariates and extra-Poisson sampling error.

Finally, we complete the model by setting priors on the parameters *β*_0_, *β, ρ* and *σ*_*u*_ and *σ*_*v*_. We assigned *ρ* and *σ*_*u*_ a joint penalised complexity prior (Fuglstad et al., 2018) such that *P* (*ρ <* 1) = 0.00001 and *P* (*σ*_*u*_ > 1) = 0.00001. This prior encoded our *a priori* preference for a simpler, smoother random field. We set this prior such that the random field could explain most of the range of the data if required.

We assigned *σ*_*v*_ a penalised complexity prior (Simpson et al., 2017) such that *P* (*σ*_*v*_ *>* 0.05) = 0.0000001. This was based on a comparison of the variance of Poisson random variables, with rates given by the number of polygon-level cases observed, and an independently derived upper and lower bound for the case counts using the approach defined in (Cibulskis et al., 2011). We found that an *iid* effect with a standard deviation of 0.05 would be able to account for the discrepancy between the assumed Poisson error and the independently derived error. Finally, we set regularising priors on the regression coefficients *β*_*i*_~ Norm(0, 0.4). The models were implemented and fitted using Template Model Builder (Kristensen et al., 2016) in R (R Core Team, 2018).

We compared the performance of the models with three sets of covariates, *X*. Firstly, we used the environmental and anthropogenic covariates, centered and standardised. Secondly, we used the predictions from the machine learning models. Finally we combined these two sets of covariates.

To compare the three models we used two cross-validation schemes. In the first, polygon incidence data was randomly split into six cross-validation folds. In the second, polygon incidence data was split spatially into three folds (via k-means clustering on the polygon centroids). This spatial cross-validation scheme is testing the models’ ability to make predictions far from data where the spatial random field is not informative. Our primary performance metric was correlation between observed and predicted data.

## 3. Results

Figure 1 shows the model performance under random and spatial cross-validation for both Madagascar and Colombia. The poor model performance in Colombia under spatial cross-validation indicates that the covariates alone cannot explain malaria incidence in this area. For all other models that use the machine learning predictions as covariates, correlations between observed and predicted data of 0.54 – 0.76 were achieved (Table 1). Input data and mapped out-of-sample predictions of the best performing model, in Colombia, are shown in in Figure 2.

**Figure 1:**
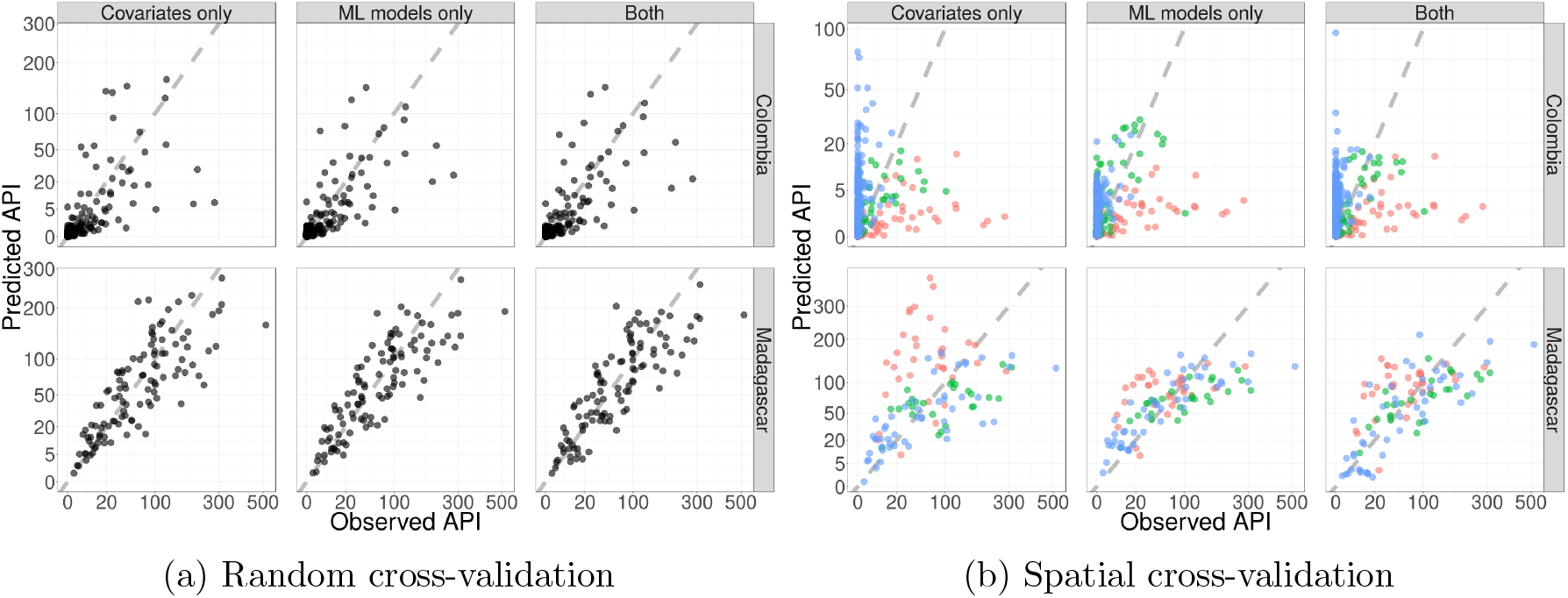
Observed data against predictions for cross-validation hold-out samples on a square root transformed scale. a) Six-fold random cross-validation. b) Three-fold spatial cross-validation with folds indicated by colour.

**Table 1:** Pearson correlations between observed and predicted values.

| Cross-validation scheme | Country | Covariates | ML | Covs + ML |
| --- | --- | --- | --- | --- |
| Random | Colombia | 0.45 | **0.55** | 0.54 |
| Random | Madagascar | 0.70 | **0.76** | 0.75 |
| Spatial | Colombia | 0.05 | **0.18** | 0.10 |
| Spatial | Madagascar | 0.22 | **0.63** | 0.61 |

**Figure 2:**
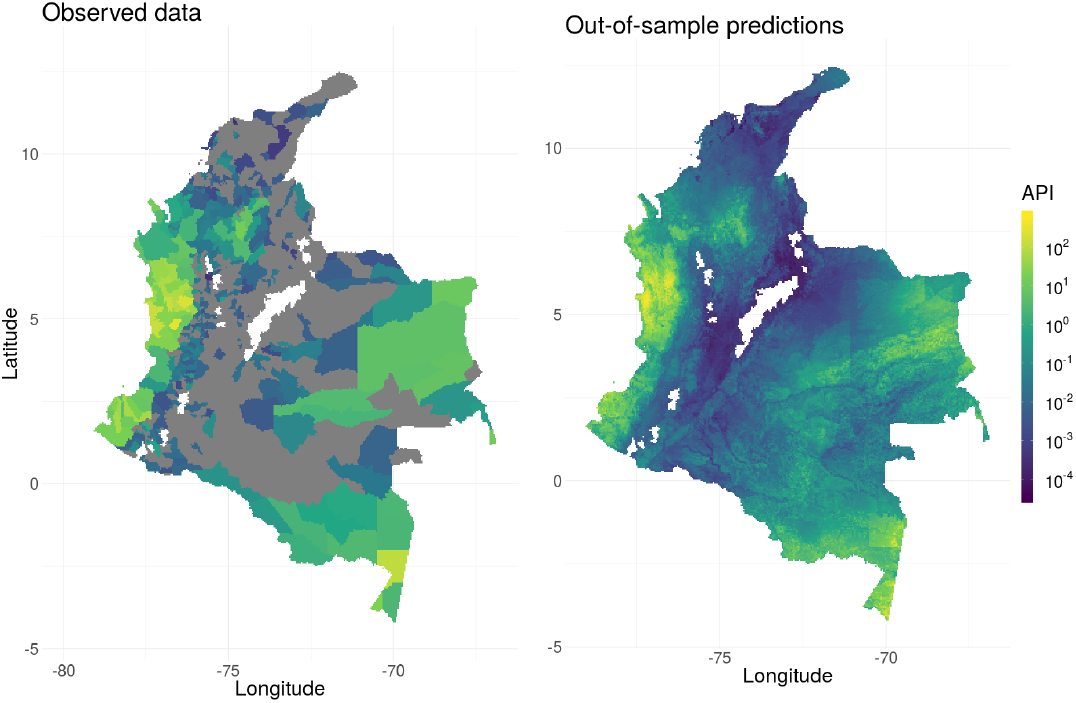
Left: Observed data for Colombia (grey for zero incidence). Right: Out-of-sample predictions for the random cross-validation, machine learning only model. For each cross-validation fold, predictions are made for the held out data which are then combined to make a single surface.

The model using only machine learning predictions as covariates was the best performing model in both countries and both cross-validation schemes (Table 1). As expected, models performed better in the random cross-validation scheme than the spatial cross-validation scheme. The difference between the covariate only model and the machine learning predictions only model was greater in the spatial cross-validation scheme than in the random cross-validation. The improvement in performance between the worst and best models was always smaller than the difference between the random and spatial cross-validation schemes.

Predictive performance of machine learning models was similar, with Random Forest performing best in Madagascar and neural networks, Random Forests and elastic net performing equally well in Colombia (Table 2). The means (across folds) of the regression coefficients (i.e. the weights of the machine learning models in the level zero model) from the polygon-level models that used only predictions from machine learning models as covariates can also be seen in Table 2. The estimated regression parameters are similar between the random and spatial cross-validation schemes. However, the best performing machine learning models do not have the largest estimated regression coefficients as would be expected if prevalence and incidence were completely correlated. Also of note is that some models were estimated to have a negative relationship with incidence (conditional on the inclusion of predictions from other machine learning models).

**Table 2:**
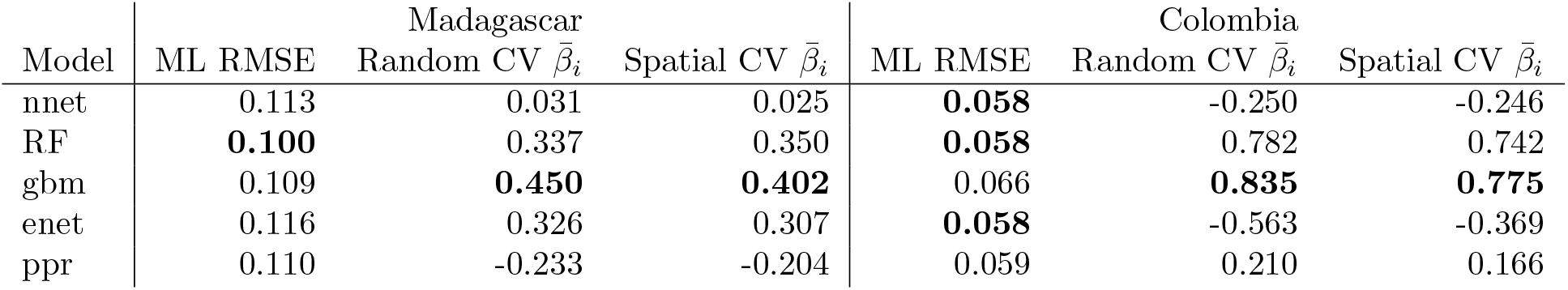
Machine learning model results and means of fitted parameters (i.e. model weights) across cross-validation folds of the machine learning predictions only model.

## 5. Conclusions

Overall, our experiments suggest that using predictions from machine learning models trained on prevalence points provides more accurate predictions than using environmental covariates when fitting disaggregation models of malaria incidence. This increased performance comes despite the data being on different scales, the data being measurements of different aspects of malaria transmission and despite the imperfect model we have used to translate between the two scales. However, the reduced model accuracy in the spatial cross-validation schemes, relative to the random cross-validation scheme, highlights that better spatial coverage of data would improve predictions more than the improved model we have suggested.

Due to the low power of typical aggregated incidence datasets, previous analyses using disaggregation regression used a small number of covariates (Sturrock et al., 2014). However, as models such as Random Forest and elastic net can robustly handle high dimensional data, future work could include many more covariates, potentially increasing predictive performance.

While the approach presented here is related to stacking, it differs in that we have not constrained the regression parameters to be positive nor included a sum-to-one constraint i.e. the result is not simply a weighted average of the level zero model predictions. We did not include these constraints because the base models and the meta model are trained on response data on different scales. However, future work could examine whether using a positive constraint on the regression parameters improves performance.

Another area of potential improvement is varying the data used to train the base level learners. Here we only used data from the region of interest. However, the global dataset is much larger than these subsets. Training some base level models on local data and some on the global dataset and then combining predictions from all these models has potential to further improve model performance.

